# Hypoxia increases the tempo of evolution in glioblastoma

**DOI:** 10.1101/293712

**Authors:** David Robert Grimes, Marnix Jansen, Robert J. Macauley, Jacob G. Scott, David Basanta

## Abstract

**Background:** Low oxygen in tumours have long been associated with poor prognosis and metastatic disease, precise reasons for which remain poorly understood. Somatic evolution drives cancer progression and treatment resistance. This process is fuelled not only by genetic and epigenetic mutation, but by selection resulting from the interactions between tumour cells, normal cells and physical microenvironment. The ecological habitat tumour cells inhabit influences evolutionary dynamics but impact on tempo of evolution is less clear.

**Methods:** We explored this complex dialogue with a combined clinical-theoretical approach. Using an agent-based-model, we simulated proliferative hierarchy under heterogeneous oxygen availability. Predictions were compared against clinical data derived from histology samples taken from glioblastoma patients, stained to elucidate areas of hypoxia / necrosis, and p53 expression heterogeneity.

**Results:** Simulation results indicate cell division in hypoxic environments is effectively upregulated, and that low-oxygen niches provide new avenues for tumour cells to spread. Analysis of human data indicates cell division isn’t decreased in low-oxygen regions, despite evidence of significant physiological stress. This is consistent with simulation, suggesting hypoxia is a crucible that effectively warping evolutionary velocity, making deleterious mutations more likely than in well-oxygenated regions.

**Conclusions:** Results suggest hypoxic regions alter evolutionary tempo, driving mutations which fuel tumour heterogeneity..

## Introduction

While genetic alterations are the fuel of cancer’s somatic evolution, the tumour micro-environment is the key contributor to the selection process that could be described as its engine^1^. The micro-environment of a tumour is made of multiple elements, all of which can have an impact on the fitness of the tumour cells, thus shaping the selection for key cancer phenotypes^1–3^ that characterize tumour progression and determine the patient’s prognosis. A key element of this micro-environment is oxygen. As early as the 1950s, investigations by Gray and colleagues^4^ showed its pivotal role in patient prognosis. Since these initial investigations, ample further evidence has emerged confirming the observation that the oxygenation of a tumour has important implications for patient outcome and treatment response^5–7^. There are two major reasons for this – the first is that poorly oxygenated tumours respond significantly worse to treatment than well-oxygenated tumour regions^8–10^; second, oxygen is a known selection pressure which favors specific aggressive cancer cell phenotypes characterized by certain known traits. Most prominent of these traits is the capacity to endure harsh environments, and the ability to migrate beyond the tissue from whence they arose^11^. Such clones gain the ability to proliferate and survive in hypoxic environments^12^. Combined, these factors suggest that hypoxia can initiate metastasis, though the exact mechanism remains unclear. For these reasons, the oxygen micro-environment is of particular interest, and has been studied extensively in physiology and pathophysiology. In healthy tissue, it is relatively stable and well-supplied. However, tumours tend to have highly heterogeneous microscopic oxygen supply^13–16^, a direct consequence of the chaotic and erratic vasculature encouraged by tumour angiogenesis^17–20^. Improving our understanding of the interplay between the oxygen micro-environment and cancer evolution is of paramount importance to advancing therapy^21–24^, yet it is notoriously difficult to probe this question with experimental tools alone.

Mathematical modeling allows us to explore the consequences of various assumptions, even when empirical observations are difficult to obtain. These models can incorporate experimental and clinical data, and the hypotheses they produce can be challenged against that coming from *in vitro, in vivo* and patient data. Where data are available, mathematical models can help inform our understanding of what is observed^25, 26^, and further, to understand the spatio-temporal dynamics to which the study of fixed tissue or molecular biology is typically blind. It has become increasingly recognized that the integration of mathematics and clinical as well as experimental data in oncology can yield novel insights that are clinically relevant^25^.

In this work, we integrate clinical observations of spatial and temporal heterogeneity in tumour oxygenation with proliferative cellular heterogeneities assuming a tumour composed of cells with different degrees of differentiation. Specifically, we developed an agent-based model, using a hybrid discrete-continuous cellular automata (HCA) approach^27^, of neutral tumour evolution in a proliferative hierarchy. Using this model, we studied the evolutionary dynamics of the clonogenic cancer cells across the heterogeneous tumour and tumour microenvironment and present evidence that hypoxic regions lead to selection for clonogenic cells and an increase in the stress of the tumour cells that inhabit it. We elucidate the implications of this, including the finding that could lead to an increase in the evolutionary tempo of these regions compared to those that are well oxygenated. This latter result was contrasted against histology from patients with this disease.

## Materials and Methods

### Clonogenic cell model outline

To explore stem cell dynamics in a heterogeneous oxygen environment, we used an agent-based HCA model built upon the framework developed previously^28^ with modification. The schematic is outlined in figure 1(a). Briefly, it consists of clonogenic cells which can symmetrically divide (with probability *α*) into two identical stem cells, or asymmetrically into a clonogenic daughter and a daughter transient amplifying cell (TAC) with probability 1 − *α*, provided there is free space for the cells to occupy. Clonogenic cells are effectively immortal unless killed by anoxia, TACs divide to other TACs only, and these cells can only undergo *β* divisions before undergoing apoptosis. TAC cell daughters inherit the divisional age of their parent TAC. An alternative explanation to our assumptions is that any cancer cell can give rise to another cancer cell. Modelling suggests that the assumptions made have serious implications for tumour growth^29^, and it is worthwhile to consider both options. To implement the assumption that all cells would proliferate, the simulation was also run with *α* = 1 so no TAC cells would emerge. To factor in the influence of the oxygen micro-environment, simulations were run with a variety of oxygen maps, with the addition of conditions for hypoxia mediated death. These maps were simulated from previously derived vascular maps / oxygen kernels^19^, scaled up to illustrate typical oxygen heterogeneity. Figure 1(b), (c), and (d) depict simulated oxygen maps derived from 1, 15, and 357 vessel configurations respectively. In regions below a critical oxygen threshold *p*_*C*_, cells have a probability *P*_*D*_ of death per time-step, simulated with both the Heaviside switch function and oxygen dependent death function, as outlined in the appendix S1. The HCA model was run considering these oxygen maps, following the evolution of cancer cells in the micro-environment, recording not only cell position but the divisional age of cells (i.e. the number of total divisions in their life history). Divisional age was taken as a proxy for mutational risk, as cells which undergo more divisions have increased chance of producing an offspring with a clinically relevant mutation (for example, conferring increased therapeutic resistance or metastatic potential). Each grid position is assumed to be the width of one-cell. For simplicity, no cellular compression was assumed. The model was run 1, 000 times over each oxygen maps outlined and output analysed. Simulation parameters are given in supplementary material S1.

**Figure 1.**
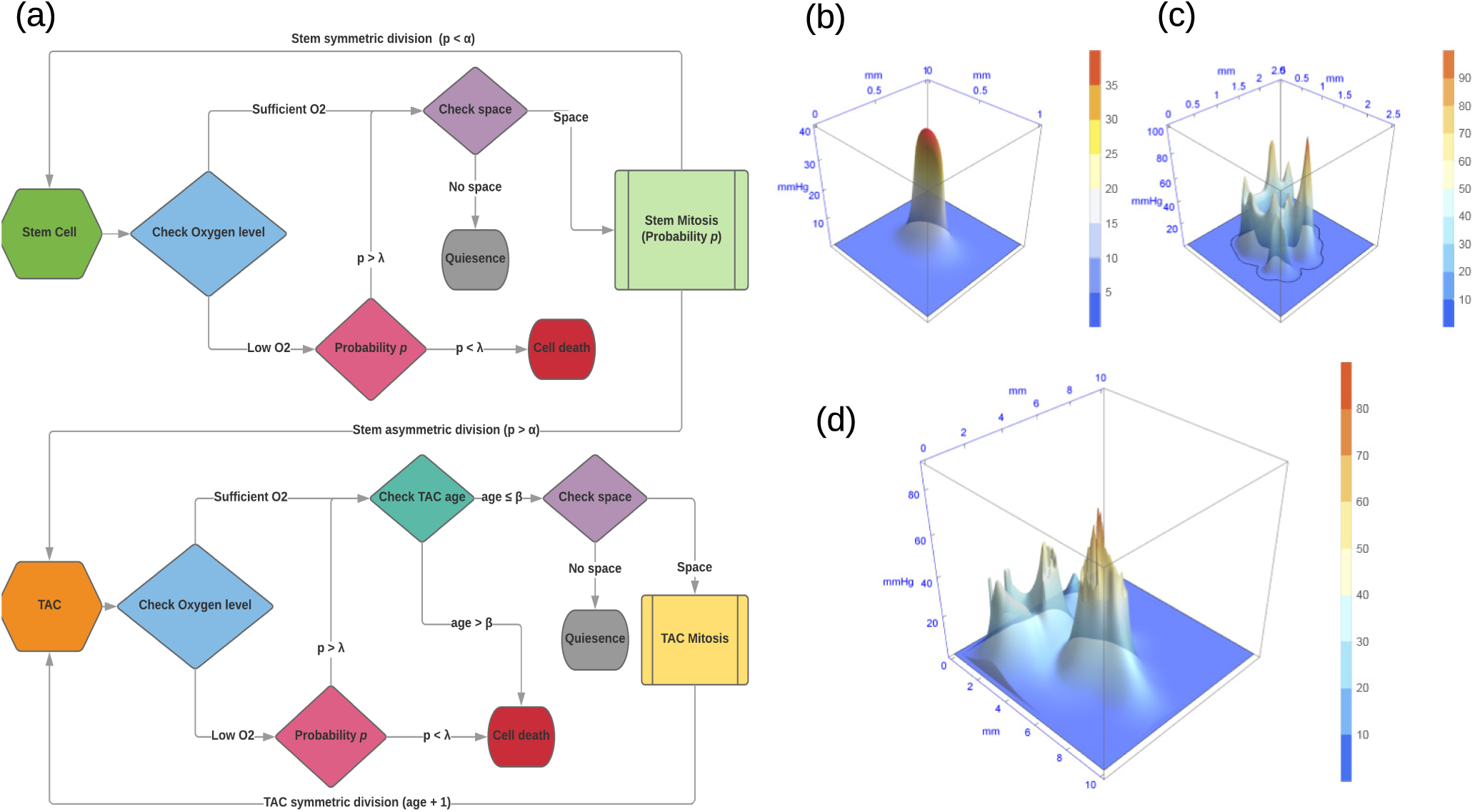
Model schematic and example oxygen maps. (a) Cell fate decisions per each cellular update are determined by the the flow charts displayed. Note that the non-italicized p in this schematic refers to probability rather than oxygen tension. Example static heterogeneous oxygen maps with (b) 1, (c) 15 and (d) 357 vessels.

### Analysis of clinical data

Human glioblastoma sections were obtained from patient biopsy samples. For each tumour, three adjacent sections were prepared as follows: 1) hematoxylin and eosin; 2) immunohistochemistry (IHC) for the proliferation marker Ki-67; and 3) IHC for p53 protein. While overexpression of the latter can sometimes be interpreted as a surrogate for TP53 gene mutation and gene dysregulation in a number of cancers^30–34^, it is chiefly an indicator of physiological cellular stress. Gene sequencing was also performed on the sections to determine whether TP53 gene mutatation was present or not, with all clinical aspects of this study approved by the Moffitt Cancer Center IRB. To quantify cellular features, microscopy was performed at high resolution using the Digital Pathology Leica Biosystems Aperio system. Images were taken at 20X magnification, yielding digital images of the sections with 1 pixel corresponding to 0.504*µ*m. Regions of necrosis were identified by histological examination on the H&E slide, and marked by a specialized neuro-pathologist (RM) using the Aperio *Imagescope* software. These annotations were extracted as XML files with the coordinates of necrotic boundaries. While explicit oxygen concentration cannot be determined from this experimental data, a major benefit of using glioblastoma sections is that necrosis in these cancers is strongly associated with hypoxia, so that necrotic boundaries could be treated as a reliable proxy for hypoxia even without explicit oxygen concentrations. This assumption is justified in more detail in the discussion section. A co-registration algorithm was written for this work, which identifies features in adjacent slides and aligns the images. Once images were co-registered, cells staining both positive and negative for Ki-67 were identified automatically on the Ki-67 slide, and P53 positive cells on the p53 slide. The image analysis code determines distance from the coordinates of each cell center to the nearest boundary of identified necrosis, recording the minimum distance to necrosis for each cell of interest. An example of the co-registration and cell identification technique are shown in figure 2. A full description of the image registration algorithm, image analysis protocol and sample code is included in the supplementary material **S1**.

**Figure 2.**
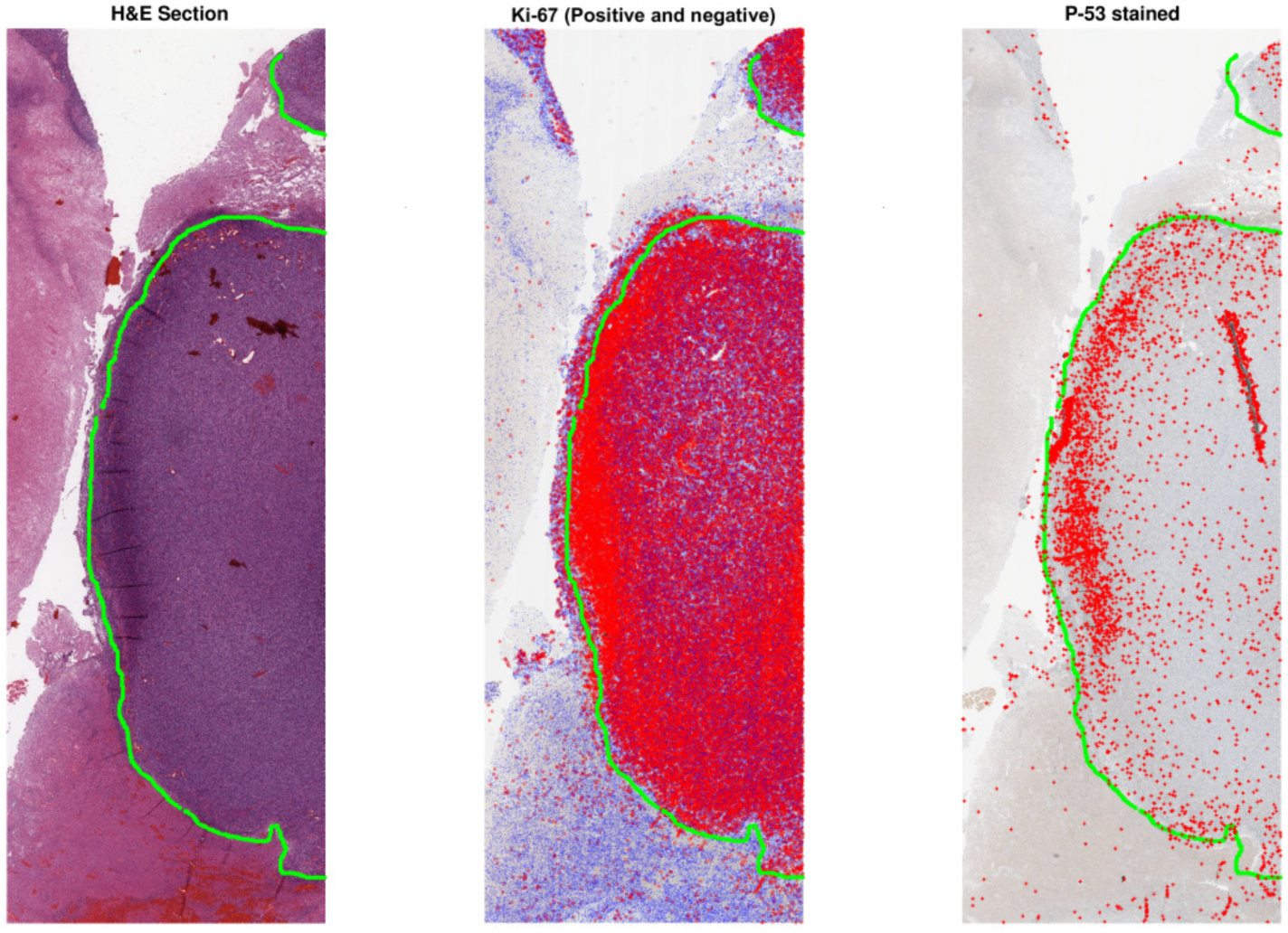
Co-registration and cell detection analysis. A necrotic boundary is marked on the the H&E slide by the pathologist (marked here by the green line). On the Ki-67 stain, cells which meet the threshold for Ki-67 positive are marked by red dots, and those below threshold by blue dots. Finally, P53 positive cells are marked by red (+) symbols on the final stain. The region shown above encompasses an area of 87.52 mm^2^ (15.67 mm × 5.58mm.)

### Evolutionary pressure of hypoxia

From the quantification of clinical data discussed above, we can now investigate the hypothesis that cells in the hypoxic niche are at higher risk of mutation. For a clonogenic cell, we assume that the rate at which mutations are accumulated per unit time, *γ* is related to the division rate *g* and the intrinsic risk of mutation per division, *r*_*d*_. It follows that for multiple divisions, *γ* = 1 − (1 − *r*_*d*_)^*g*^. When *r*_*d*_ is small, the binomial approximation allows us to *γ* ≈ *r*_*d*_*g*, and thus the probability of a clonogenic cell acquiring a mutation with time *t* is given by Poisson statistics as *M*(*t*) = 1 − exp(−*r*_*d*_*gt*). Under conditions of high cellular stress, as those in the hypoxic niche, we can expect a higher intrinsic probability of mutation per division *r*_*s*_, where *r*_*s*_ > *r*_*d*_, reflecting the evolutionary pressures of the microenvironment on cellular evolution^35^. We define the mitotic rate in the hypoxic niche as *g*_*s*_ and thus

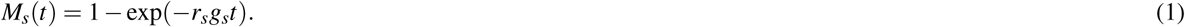

If hypoxia leads to an increase in mutation rates, then we would expect *P*_*s*_(*t*) > *P*(*t*), ∀*t* > 0 where *P*_*s*_(*t*) is the probability of a mutation under hypoxia and *P*(*t*) the probability of mutation under normoxia. Determining this requires us to probe the mitotic status of the hypoxic niche. There is evidence that cells in the hypoxic niche respond to stress by entering a state of quiescence^36–38^, markedly reducing their rate of mitosis (*g*_*s*_ << *g*). In this case it is possible that *M*_*s*_(*t*) < *M*(*t*), which would imply hypoxia is not a selection pressure for evolutionary change. Alternatively, if there is evidence that cells in the hypoxic niche continue to undergo the same approximate rate of mitosis as cells in well-oxygenated regions, then it follows that mutations will be much more likely to arise in hypoxic niches. There is good biological evidence that hypoxia diminishes DNA repair and elevates mutagenesis^39^. Using the histological analysis outlined, the distribution of both p53 positive (physiologically stressed) cells and mitotically active Ki-67 positive cells were quantified in different regions to determine whether mutational risk was elevated under hypoxia, and results contrasted with model predictions.

## Results

### Model-derived results

#### Oxygen-dependent distribution of clonogenic cells

Figure 3 depicts the stratification of clonogenic cells relative to oxygen concentration. The vertical axis in a(i), b(i) and c(i) has been normalized to maximum number of simulated divisions in each case and presented as a ratio to faciliate ease of comparison. What is immediately apparent in simulation is that high division of clonogenic cells was directly associated with low oxygen conditions. This trend was seen for every map, regardless of the underlying heterogeneity. For all configurations, clonogenic cells on the anoxic border underwent far more divisions than well-oxygenated cells, as illustrated in Figure 3. Qualitative observation of the HCA reveals that this increase in divisional age is secondary to cyclic instances of birth and death as cells place daughters into areas of extreme (lethal) hypoxia. So while the daughters are dying, the clonogenic cells continue to divide as they sense free space. With *α* = 1 (assuming all cells clonogenic) the same trend was observed, with cells on the anoxic boundary undergoing far more divisions than those in well-oxygenated regions. Figure 4 illustrates the dynamics over a very simple oxygen map, suggesting strongly that at the hypoxic niche, divisional age of clonogenic cells is markedly upregulated, increasing mutational risk.

**Figure 3.**
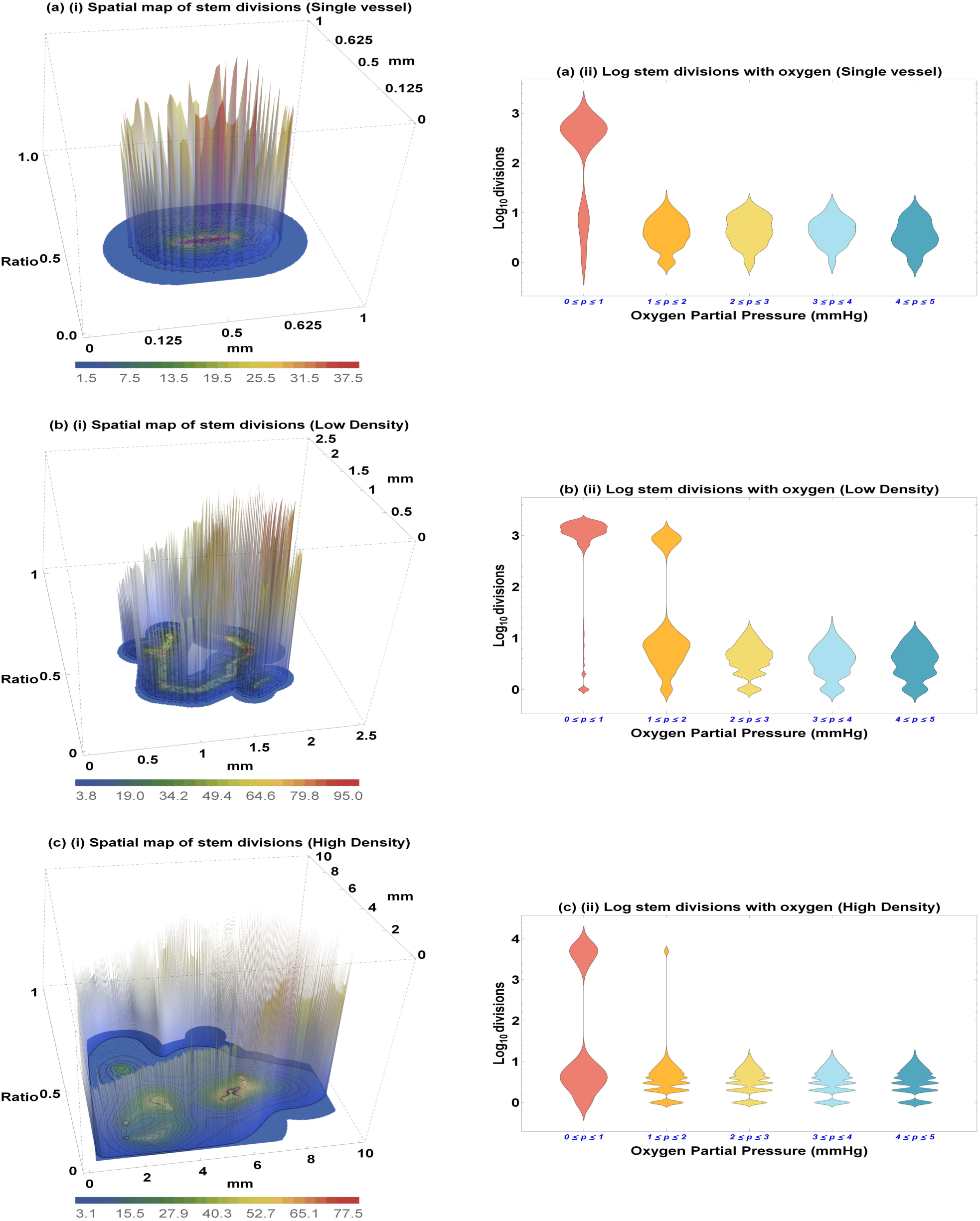
Clonogenic division with varying oxygen concentrations in (a)(i) single vessel oxygen map b(i) low density 15 vessel oxygen map (c)(i) high density 357 vessel oxygen map. Color bars indicate oxygen partial pressure, and is plotted on the horizonal plane of the figures. Divisions of all clonogenic cells are normalized to the maximum number of divisions in the vertical axis for clarity. Figures (a)(ii), (b)(ii), and (c)(ii) show clonogenic cell divisions to be higher under hypoxia.

**Figure 4.**
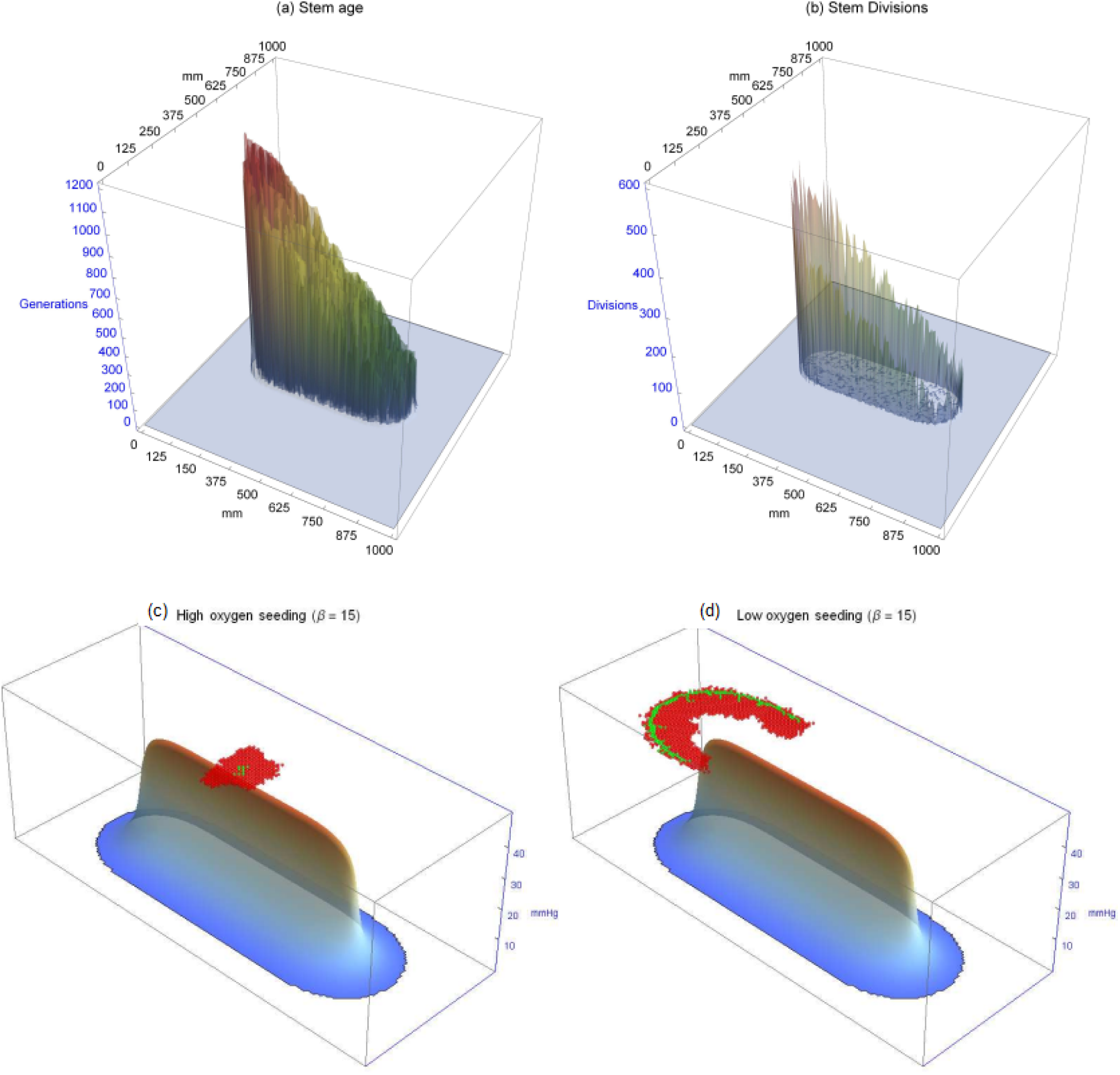
(a) Relative ages of clonogenic cells. (b) Number of clonogenic divisions for the same cells. This suggests that number of clonogenic cell divisions is correlated chiefly with low oxygen tension rather than clonogenic cell age. At the anoxic border, mitosis is markedly up-regulated. For this simulation, a vessel-like oxygen map was used with *β* = 4 and *t* = 1, 200. All other parameter values are as per supplementary material S1. (c) and (d) depict clonogenic cells (Green) and TACs (Red) with *β* = 15 after 10,000 time-steps for (c) clonogenic cells initially seeded in high oxygen environment (d) initial cell seeded on hypoxic niche. In (c) the fire-wall effect of long-lived TACs can be clearly seen whereas in (d) clonogenic cells can proliferate along the anoxic ridge, yielding an ‘edge creep’ effect which allows clonogenic cells to invade along hypoxic ridge.All other parameter values are as per supplementary material S1.

#### Hypoxic niche as a metastatic avenue

Figure 4 (c) and (d) depicts the impact of seeding an initial clonogenic cell in either an oxygen-rich or hypoxic environment. For high values of *β*, a clonogenic cell seeded in high oxygen yields a ‘fire-wall’ effect, where long-lived TACs impede the invasion potential, previously observed by other authors^40,41^. However, results from these simulations suggest this fire-wall is overcome when clonogenic cells are on a hypoxic border. In this instance, cells can ‘creep’ along the edges, travelling along the hypoxic niche as depicted in 4(d). This leads to marked differences in the clonogenic population; for the simulation described, seeding in high oxygen with *β* = 15 led to only 13 ± 4 stem cells after 10,000 steps. By contrast, seeding in low oxygen yielded 254 ± 25 clonogenic cells after the same time had elapsed.

### Clinical data analysis

Clear borders of necrosis could be ascertained in 23 sections from 9 patients, details of which are given in the supplementary material S1. These sections for analysis ranged from 0.72mm^2^ to 108.14mm^2^. For each section, image analysis was performed to determine cells that were both positive and negative for Ki-67, and for cells positive for P53 mutations. With these cells and their positions determined, the minimum distance from the cell to the pathologist-specified necrotic boundary was calculated. The probability density of spatial distribution from known necrosis for all this data is shown in Figure 5, in bins corresponding to the width of two cells (25 *µ*m). There was no statistical difference in the distribution of cells both positive and negative for Ki-67 relative to necrosis (two-sample Kolmogorov-Smirnov test *p* = 0.5668, KS test statistic 0.0802), and accordingly these are grouped together. By contrast, P53 mutation-positive cells are far more likely to be found near regions of necrosis, and have a markedly different distribution than the grouped Ki-67 cells (two-sample Kolmogorov-Smirnov test *p* = 1.21 × 10^−7^, KS test statistic 0.2941). Gene sequencing did not show any indication of TP53 mutation, which strongly implies that the P53+ stained cells were indicative of physiological stress, likely hypoxia-driven.

**Figure 5.**
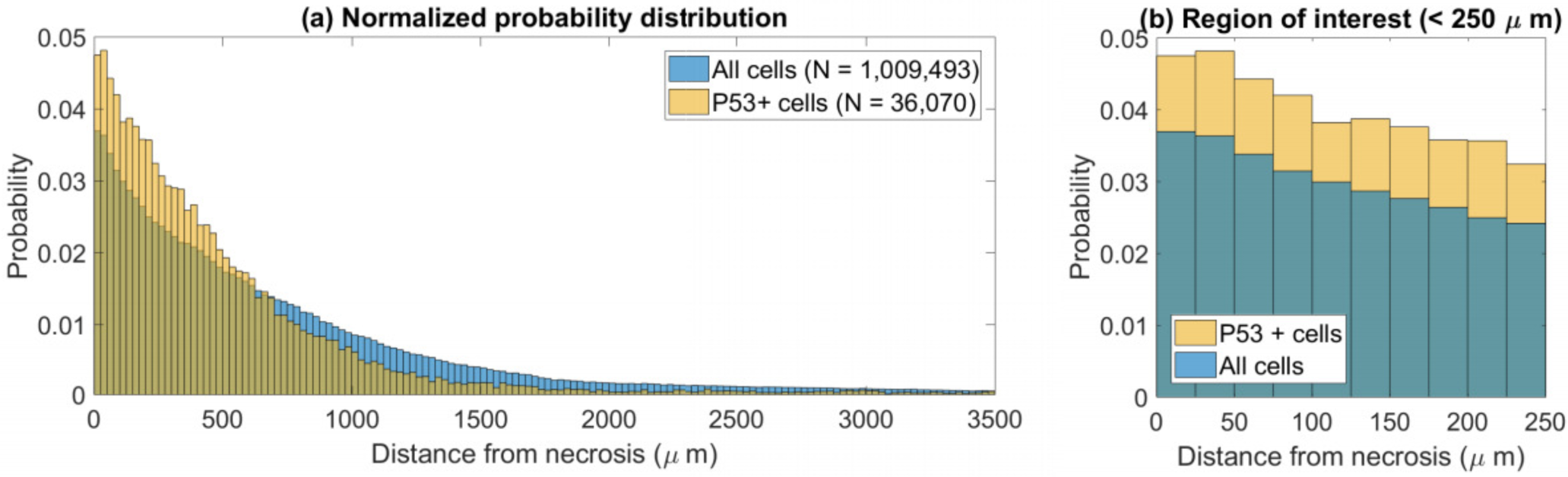
Physiological stress more probable close to known regions of necrosis. Pooled data from 23 regions of 9 patient glioblastoma samples after image analysis. (a) Distribution of P53 stained cells versus Ki-67 stained cells relative to known necrotic borders. (b) Probability distributions for stained cells close to necrosis.

#### Analysis of evolutionary risk

To determine whether cells in the hypoxic niche displayed greater quiesence, the fraction of Ki-67 positive cells over the sum of Ki-67 positive and negative cells in each 25 *µ*m bin from 0-2mm bin was calculated. This fraction was calculated at 0.2124 ±0.0104, indicating that the proportion of mitotic cells in each bin was relatively constant, and that *g*_*s*_ ≈ *g*. By contrast, p53 staining was markedly increased close to regions of hypoxia (see supplementary S1 for correlation data), suggesting strongly that cells under physiological stress continued to undergo unrestricted mitosis. This suggests that cells in the peri-necrotic niche have increased mutational risk relative to well-oxygenated cells.

#### Hypoxia and evolutionary tempo

The clinical data described above shows an increase in physiological stress associated with hypoxia. This is illustrated clearly in figure 6, which depicts a histological section stained with H&E. In the figure, cells straining strongly positive for p53 mutation as detected by the image analysis in the p53 section are superimposed at their corresponding positions, marked by blue dots. Regions of clear necrosis as demarcated by the neuropathologist (RM) are outlined in green on this image. From the spatial map of p53 mutant cells, a probability distribution function for these points in space was ascertained by employing a Sheather-Jones data smoothing kernel in *Mathematica*, which yields the non-parametric probability density function of a random variable^42^. From this, contour lines of p53 stress density have been superimposed over the image, with greater line density denoting increased abundance of p53 staining cells. For ease of viewing, a red opacity effect has also been superimposed over the image to show the highest density of p53 staining cells, under physiological stress. This corresponds to the contour lines of concentration. As can be seen, the highest density of stressed cells tend to lie on or close to the green line of representing the necrotic anoxic boundary, illustrating stress near the necrotic hypoxic boundaries. This echoes the phenomena predicted in our simulations, with the resultant map yielding the likely topography of evolutionary velocity.

**Figure 6.**
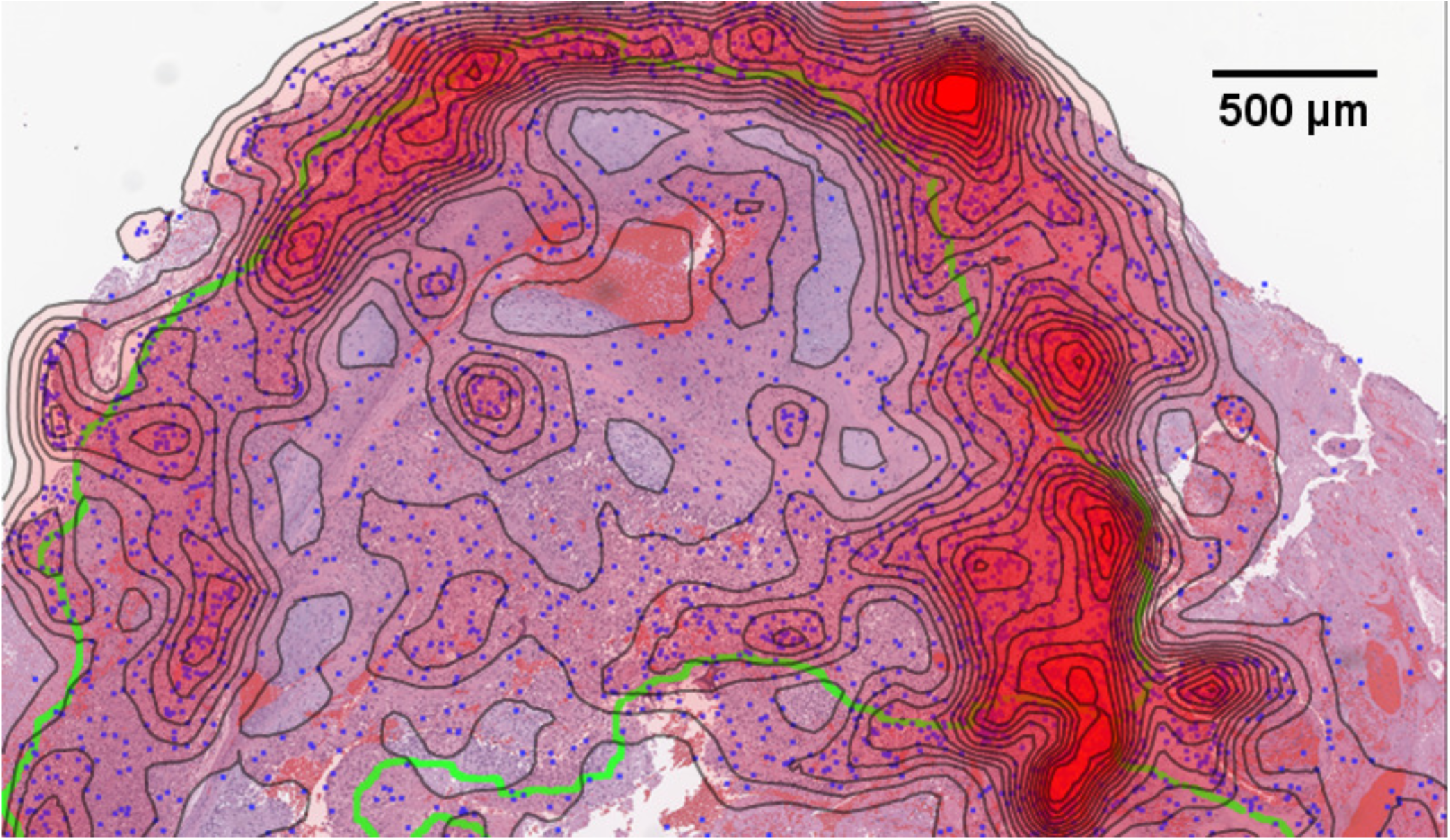
Physiological stress indicates potential topography of evolutionary velocity. An example from a patient glioblastoma histologic section. Physiological p53 stress detected by image analysis is illustrated by blue dots overlaid on the histology section. Green lines depict pathologist marked necrosis, and contour lines with red opacity show the probability density of stress markers (calculated from a Sheather-Jones smooth kernel distribution function). Near necrotic regions, the probability of finding stress markers increases relative to non-necrotic zones. As mitotic rate appears approximately constant throughout tissue, this suggests these regions are more likely to give rise to mutations.

## Discussion

Hypoxia and necrosis are known hallmarks of cancer, and literature to date indicates that hypoxia selects for aggressive and metastatic phenotypes. In this work we have investigated the hypothesis that hypoxia can also influence the speed and evolutionary potential of cancer, acting as a potential strong selection pressure for subclonal evolution as defined in recent works by other researchers^43^. We further present evidence that this impact goes beyond selecting for certain phenotypes more adaptable to low oxygen levels, but that hypoxia directly modulates the tempo of somatic evolution. Specifically, we show that the field representing the speed of somatic evolution is significantly increased near the anoxic edges surrounding areas of necrosis. To draw an analogy from physics, the presence of hypoxia appears to ‘warp’ evolutionary velocity – effectively creating a regions of increased rate of potential mutation acquisition, which are micro-environmentally mediated evolutionary hot-spots.

To come to these conclusions, we developed and interrogated an HCA computational model and found that clonogenic cell division is substantially more pronounced in regions of hypoxia relative to more well oxygenated regions, as illustrated in figure 4. The simulations also revealed that the hypoxic niche can facilitate the migration of clonogenic cells along low oxygen regions, markedly increasing their population. Other authors have found that long-lived TAC cells effectively limit cancer growth, acting as a firewall when these offspring cells are sufficiently long-lived^40^. This model recapitulated that behaviour in well-oxygenated environments, but this behaviour broke down in the presence of an anoxic niche, as illustrated in figure 4, where clonogenic cells were able to colonize the necrotic niche, despite TAC cells in this simulation being very long lived (*β* = 15). This ‘edge effect’ suggests that the hypoxic niche can act as a conduit to cellular infiltration, effectively changing the way cells interact^44^. Whilst further biological evidence will be required to confirm this, it raises a previously unforeseen potential consequence of hypoxia for tumour evolution.

Histopathological data from glioblastoma patients was examined to challenge *in silico* predictions, and to determine whether modeled behaviour was consistent with it. We performed image-analysis on sectioned regions from glioblastoma patients, with co-stained sections. This strongly suggests that cells in the hypoxic niche do not undergo any noticeable quiesence, displaying the same fraction of Ki-67 proliferation marker as well-oxygenated regions. This was consistently observed, including in areas with clear markers of severe physiological stress. The analysis in this work suggests that tumour cells continue to proliferate unimpeded by the stressful conditions they find themselves in, increasing their risk of mutation. This isn’t unprecedented - insensitivity to signaling and persistent proliferation are of course hallmarks of cancer^45^. There is ample evidence that hypoxia elevates mutagenesis^39, 46, 47^, and modeling results in this work suggest a mechanistic reason why cells in the hypoxic niche would be far more likely to acquire mutations than well-oxygenated cells, leading to eventual emergence of metastatic or treatment resistant types. Increased division by clonogenic cells in the hypoxic niche elevates the probability of a cell acquiring a mutation (and ultimately metastatic potential) which can in part explain why hypoxia is so strongly correlated with the emergence of metastatic phenotypes and poor prognosis^11^. The biological evidence here is of course indirect due to the limitations of staining analysis, and dedicated and specialized experiments would be needed to fully test the hypothesis fully. Importantly however, observed data is in accordance with simulation predictions based on mechanistic principles, suggesting that this is a fertile avenue for future exploration.

The model presented in this work is a simple agent-based model which was kept as general as possible. We do not expect it to capture all the complexities of a true biological system, but rather to approximate likely behaviours which could emerge in different cancers. In the model presented, cells can either be killed off in the hypoxic zones or in the case of TACs, undergo apoptosis after *β* divisions. This prompts the question of whether small amounts of random death might change the trends observed. To test this, the simulations were also run with random death. For biologically reasonable estimates, results were similar with that presented here, illustrated in supplementary material **S1**. There are of course a number of limitations to our modeling approach, chiefly that the model exists on a simple 2D grid rather than true 3D space, and that cell motility was not considered. Increased detail and consideration of such factors could provide how precise the model is, but we do not expect the main conclusions to be challenged in light of these considerations.

On the theme of histologic evidence, a number of caveats have to be kept in mind when interpreting such data. Primarily, 2D histology is at best an approximation of complex 3D behaviour, and can in some instances be misleading^19^. Another confounding factor was the hardship of defining necrosis robustly - while one of us is a trained neuro-pathologist capable of demarcating clear regions of necrosis, there were sections which were ambiguous and were left out of the analysis for that reason. This means that the extent of necrosis may in some instances be an underestimate. Even so, a number of suitable sections were unambiguously identified in the patient data, amounting to over a million individual cells. With such a volume of data, we expect general patterns to become apparent even with the confounding influence of 2D data.

Interpretation of the clinical data pivots on the implicit assumption that regions of necrosis marked by the clinician were hypoxic. One major attraction of using glioblastoma sections is that there is ample evidence that regions of necrosis are hypoxic^48–53^. Pseudopalisading necrotic cells in particular are known to be hypoxic, displaying dramatic up-regulation of hypoxia inducible factor-1^53^. CA-IX immunostaining was also performed on some of the cases in this work, which confirmed the perinectroic regions were indeed hypoxic. That oxygen-diffusion limited hypoxia gives rise to necrosis has long been observed in human tissue^54^ and experimental models^55^, and there is a known reciprocal relationship between p53 and hypoxic path^56^. Necrotic borders in this work are almost certainly hypoxic, but for future investigations, the ability to quantify the oxygen gradient may yield further insight into the implications for tumour evolution.

This work presents combined modeling and experimental evidence that the oxygen micro-environment plays a fundamental role in ‘warping’ the evolutionary velocity of cells under its influence. This highlights the importance of the tumour microenvironment not only in selecting for certain phenotypes but also in regards to the velocity and dynamics characterizing its somatic evolution. Hypoxia itself is already detrimental for treatment efficacy^5, 8^ and this work further suggests this could be compounded by the ability of this environment to select for phenotypes displaying both increased treatment resistance, evolutionary, and metastatic potential. This suggests that hypoxic zones are of substantial pathological interest in terms of tumour evolution, and be a fruitful avenue for future investigations.

## Supporting information

Updated S1 file

## Funding statement

DRG thanks Cancer Research UK for the travel grant that made this work possible, and the Wellcome trust for their support. The authors would also like to thank the Integrated Mathematical Oncology department at the H. Lee Moffitt Cancer Center and Research Institute. DRG also acknowledges the contributions of NVIDIA research for their generous hardware donations whilst DB acknowledges the National Institute of Cancer (NCI) for grant U01CA202958-01. JGS is grateful to the NIH Loan Repayment program, the NIH Case Comprehensive Cancer Center (support grant P30CA043703), and the Calabresi Clinical Oncology Research Program, National Cancer Institute (award number K12CA076917). The funders had no role in study design, data collection and analysis, decision to publish, or preparation of the manuscript.

## Notes

#### Summary of Updates

Highly revised version, with new data and clarification of older ambiguous sections, and updated author lists. Supplemental data also updated

