## Supplementary material for "Hypoxia increases the tempo of evolution in glioblastoma": Updated S1 file

### Hypoxia increases the tempo of evolution in glioblastoma - Supplementary material S1

#### ABSTRACT

Supplementary material **S1** for paper.

#### Patient sample data

Table 1 lists details of the patient sample data used in this work, including the number of cells of each type identified in the section. Figure 1 shows clear correlation between distance from necrotic borders and probability of p53 mutation staining, suggesting physiological stress in these regions consistent with sustained hypoxia.

**Table 1.** Analysis of experimental Glioblastoma sections

| Patient | Sub-section | Area | Ki-67 + | Ki-67 - | P53 + | TP53 mutation |
|---|---|---|---|---|---|---|
| 1 | i | 108.14 $mm^2$ | 57,498 | 208,789 | 4217 | - |
| | ii | 87.42 $mm^2$ | 53,435 | 132,068 | 3991 | - |
| 2 | i | 27.21 $mm^2$ | 13,709 | 63,052 | 3072 | - |
| | ii | 21.10 $mm^2$ | 5814 | 57,689 | 1516 | - |
| 3 | i | 25.82 $mm^2$ | 10,419 | 55,213 | 2722 | - |
| | ii | 10.08 $mm^2$ | 4794 | 20,108 | 1983 | - |
| | iii | 18.62 $mm^2$ | 13,580 | 33,186 | 4547 | - |
| | iv | 15.83 $mm^2$ | 5090 | 53,798 | 275 | - |
| | v | 4.93 $mm^2$ | 2249 | 14,333 | 120 | - |
| 4 | i | 4.09 $mm^2$ | 2028 | 9569 | 360 | - |
| 5 | i | 2.17 $mm^2$ | 1956 | 4349 | 1676 | - |
| 6 | i | 2.61 $mm^2$ | 1085 | 6125 | 37 | - |
| | ii | 1.27 $mm^2$ | 500 | 3039 | 33 | - |
| | iii | 5.18 $mm^2$ | 2736 | 12,905 | 179 | - |
| | iv | 0.72 $mm^2$ | 385 | 1935 | 7 | - |
| | v | 2.89 $mm^2$ | 943 | 5985 | 252 | - |
| 7 | i | 16.10 $mm^2$ | 14,072 | 39,848 | 4373 | - |
| 8 | i | 10.88 $mm^2$ | 4275 | 10,845 | 1659 | - |
| | ii | 26.77 $mm^2$ | 14,406 | 17,757 | 4600 | - |
| | iii | 4.47 $mm^2$ | 2565 | 11,025 | 164 | - |
| 9 | i | 3.98 $mm^2$ | 1719 | 8090 | 92 | - |
| | ii | 3.79 $mm^2$ | 1810 | 7930 | 149 | - |
| | iii | 4.52 $mm^2$ | 1566 | 14,490 | 46 | - |

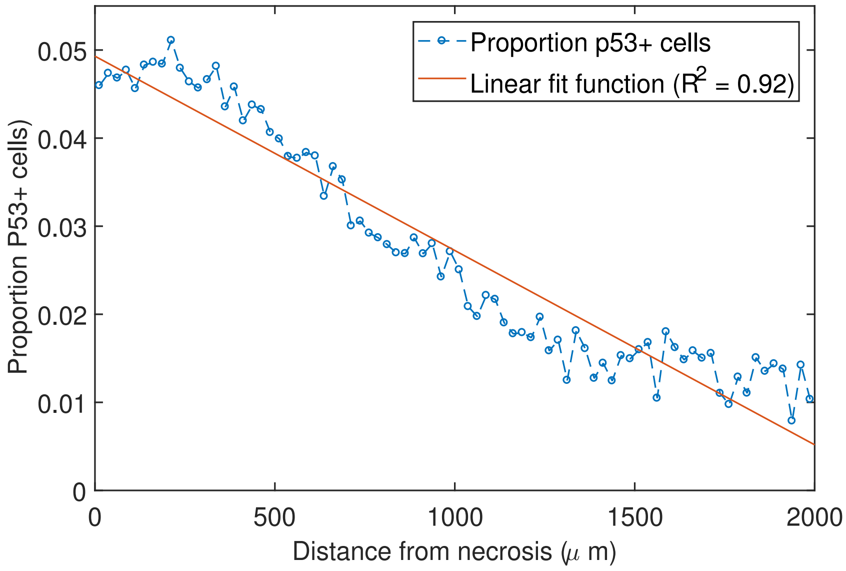


**Figure 1. Evidence of Strong physiological stress near necrotic regions** By contrast, mitotic fraction did not vary with distance from necrosis.

#### Modeling considerations

##### Guide to assumptions and equations

- Our model assumes a heterogeneous tumor made of two subpopulations: clonogenic cells capable of tumor initiation and differentiation (CSCs) and transit amplifying cells (TACs).
- CSCs can replicate indefinitely while TACs have a finite replicative potential of $\beta$ divisions after which cells die.
- CSCs can divide either symmetrically (with probability $\alpha$) or asymmetrically (where the two daughter cells will be CSC and TAC, with probability $1-\alpha$). TACs only divide symmetrically resulting in two TACs.
- Space is discretized into a grids up to 1000 x 1000. Time is also discretized into timesteps of one average cell doubling time.
- The micro-environment is determined by heterogeneous oxygen maps derived from our previous work[?].
- Cells (both CSC and TAC) in low oxygen grid points ($p \leq p_C$) have probability $P_D(p)$ of dying on every time-step. We also assume that proliferation is not impacted by $O_2$ supply.
- Cells (both CSC and TAC) divide any time that space is available and remain quiescent when there is none.

##### Key equations

Oxygen maps were derived from a previously published oxygen kernel for vascular maps, where partial pressure $p$ at a distance $d$ from a vascular point of radius $r_o$ is given by

$$p = \frac{a\Omega s_L}{3D}\left(\frac{\sqrt{r_n^2 - d^2}}{3}(2r_o^2 - 8r_n^2) + 2r_n^3 \log\left(\frac{r_n - \sqrt{r_n - d^2}}{d}\right)\right) \tag{1}$$

where $D$ is oxygen diffusion constant in water, $r_n$ is the diffusion distance of oxygen in a specific tissue and $\Omega$ and $s_L$ are constants, as previously outlined and omitted here for brevity[?]. The probability ($P_D$) of a cell dying in a low oxygen niche ($p \leq 0.5$mmHg) was modeled by two methods. The first of which was a Heaviside step function, so that

$$P_D = \begin{cases} 0.5, & p \leq 0.5 \text{ mmHg} \\ 0, & p > 0.5 \text{ mmHg}. \end{cases} \tag{2}$$

The second form allows us to capture the possibility that death probability is dependent on oxygen partial pressure, we employed a Poisson-like death function of

$$P_D(p) = 1 - \exp(-k_D p) \tag{3}$$

where $k_D$ is a constant.

##### Additional parameter table for simulations

**Table 2.** Simulation parameters

| Parameter | Value |
|---|---|
| Symmetric division probability $\alpha$ | 0.25 |
| Asymmetric division probability $(1-\alpha)$ | 0.75 |
| TAC replications before apoptosis $\beta$ | 6[?] |
| Critical oxygen threshold $p_C$ | 0.5 mmHg[?,?] |
| Cell diameter | 12.5 $\mu$m[?,?] |
| Poisson-like death function constant $k_D$ | 1.368 mmHg $^{-1}$ † |

†Chosen so that $P_D(0.5 \text{ mmHg}) = 0.5$

#### Random cell death

In the main paper, model results were presented assuming a 'no-random-death' assumption. In this framework, stem cells were effectively immortal and could only die under hypoxic conditions. TAC cells could die under hypoxia, or after undergoing $\beta$ divisions. It is worthwhile to check whether the results observed in the modeling work hold if random cell death of both stems and TACs is factored in. Supplementary figure 2 shows the impact of this for random death per cell per time step $r_d$ of 0.02 and 0.05 respectively in the low density rat tumor for 5000 time steps. In the former case ($r_d = 0.02$), results are very similar to those in the main work. In the latter case, the general trend is seen albeit with much reduced probability. This is because this high biologically improbable value for $r_d$ has a tendency to wipe out colonies; the chances $S$ of a cell surviving for $n$ iterations is $S = (1 - r_d)^n$. If $r_d = 0.02$, then a cell can live through $n = 149$ time steps before its survival chances falls to below 5%. By contrast, when $r_d = 0.05$, a cell has less than 5% chance of survival by 59 time steps. Even with these exaggerated dynamics, highly replicating stem cells were still much more likely on the necrotic niche, suggesting modeling results are robust.

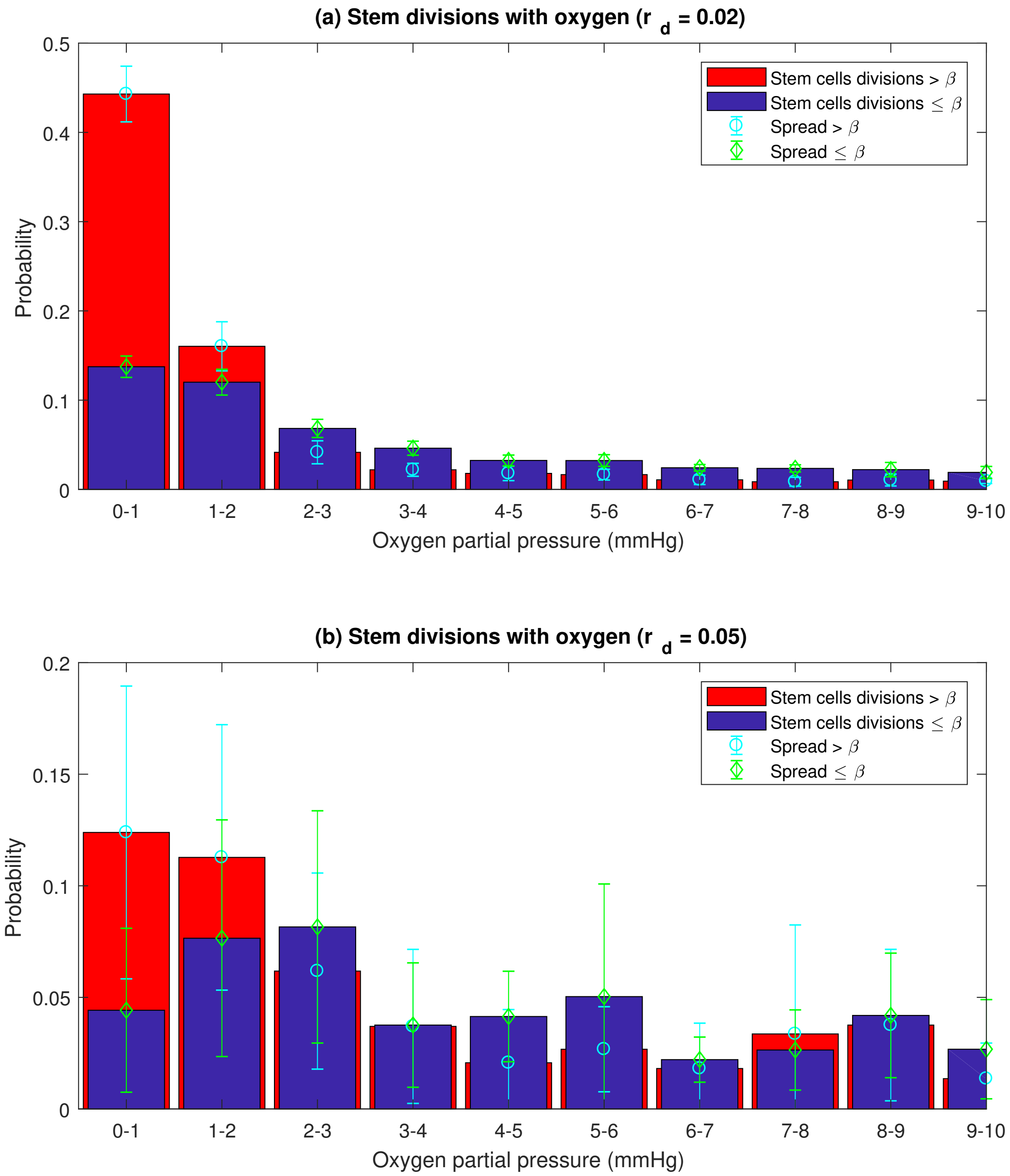


**Figure 2.** Stem cell distribution with oxygen with random death in model.

#### Image thresholding

For p53 analysis, it was important to sufficiently threshold the image so that only unequivocal cells were observed. This was to discriminate against clear mutation (strong staining) and potential physiological up-regulation (weak staining). The following algorithm was used to determine a punishing threshold, and apply it to p53 images.

- Read in red channel of p53 image.
- Convert image to grid of doubles (converts pixel values to values between 0 and 1).
- Invert image so P53 spots have high intensity.
- Find the mean pixel intensity in image, $m_{val}$.
- To reduce false positives, set threshold to a multiple $n$ of $m_{val}$, where $n > 1$.
- Binarize the image to this threshold.
- Clear any border pixels.
- Remove any small objects of less than 70 pixels.
- Find centroids of remaining objects.
- Draw over original image.
- Visually inspect, adjust $n$ as required.

For our p53 images, $n = 3$ was sufficiently high to gate ambiguous cells. This punishing cut-off might have meant that some p53 mutations were under counted, but even using lower thresholds (such as n = 2.5) yielded the same trend, provided $n >> 1$. An illustration is provided in figure 3. As this threshold was quite punitive, it should only select for the most unequivocally staining cells. This suggests weaker physiological up-regulation isn't skewing analysis.

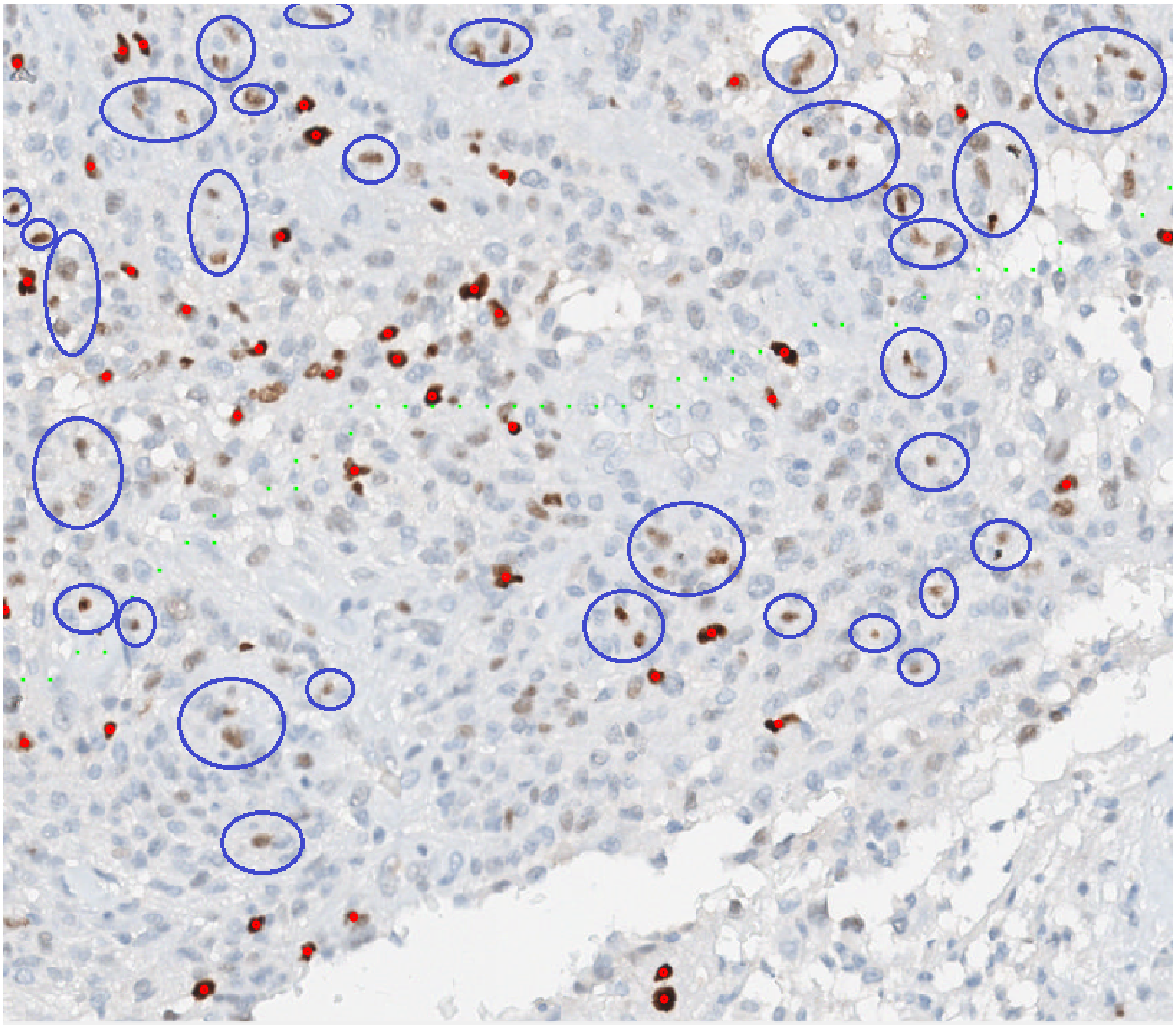

**Figure 3.** A small section of a large tumor sample. Cells beyond the threshold are shown with a small red circle. Ambiguous cells are circled in blue for illustration. The algorithm used for image analysis only selected the most unequivocally expressing cells.
